# Sophisticated suction organs from insects living in raging torrents: Morphology and ultrastructure of the attachment devices of net-winged midge larvae (Diptera: Blephariceridae)

**DOI:** 10.1101/666537

**Authors:** Victor Kang, Richard Johnston, Thomas van de Kamp, Tomáš Faragó, Walter Federle

**Affiliations:** Department of Zoology, University of Cambridge, Cambridge, United Kingdom; College of Engineering, Swansea University, Wales, United Kingdom; Laboratory for Applications of Synchrotron Radiation (LAS), Karlsruhe Institute of Technology (KIT), Kaiserstr. 12, 76131 Karlsruhe, Germany; Institute for Photon Science and Synchrotron Radiation (IPS), Karlsruhe Institute of Technology (KIT), Hermann-von-Helmholtz-Platz 1, 76344 Eggenstein-Leopoldshafen, Germany

**Keywords:** Suction organ, biomechanics, morphology, micro-CT, microstructures, Blephariceridae, adhesion, wet adhesion

## Abstract

Suction organs provide powerful yet dynamic attachments for many aquatic animals, including octopus, squid, remora, and clingfish. While the functional morphology of suction organs from various cephalopods and fishes has been investigated in detail, there are only few studies on such attachment devices in insects. Here we characterise the morphology, ultrastructure, and *in vivo* movements of the suction attachment organs of net-winged midge larvae (genus *Liponeura*) – aquatic insects that live on rocks in rapid alpine waterways where flow rates can reach 3 m s^-1^ – using scanning electron microscopy, laser confocal scanning microscopy, and X-ray computed micro-tomography (micro-CT). We identified structural adaptations important for the function of the suction attachment organs from *L. cinerascens* and *L. cordata*. First, a dense array of spine-like microtrichia covering each suction disc comes into contact with the substrate upon attachment. Similar hairy structures have been found on the contact zones of suction organs from octopus, clingfish, and remora fish. These structures are thought to contribute to the seal and to provide increased shear force resistance in high-drag environments. Second, specialised rim microtrichia at the suction disc periphery form a continuous ring in close contact with a surface and may serve as a seal on a variety of surfaces. Third, a V-shaped cut on the suction disc (the V-notch) is actively peeled open via two cuticular apodemes inserting into its flanks. The apodemes are attached to dedicated V-notch opening muscles, thereby providing a unique detachment mechanism. The complex cuticular design of the suction organs, along with specialised muscles that attach to them, allows blepharicerid larvae to generate powerful attachments which can withstand strong hydrodynamic forces and quickly detach for locomotion. Our findings could be applied to bio-inspired attachment devices that perform well on a wide range of surfaces.

## 4. Background

Firmly attached to rocks in rapid alpine streams, rivers, and bases of waterfalls, the aquatic larvae of net-winged midges (Diptera: Blephariceridae) have fascinated entomologists for over a century due to their powerful adhesion and complex attachment organs [1–4]. In these challenging environments, where the water velocity can reach up to 3 m s^-1^ [5], the midge larvae spend most of their development cycle attached to underwater rock surfaces, where they graze on diatoms from the rocks and eventually settle to pupate (Fig. 1; supplementary video SV1). Such tumultuous conditions are favourable for these larvae as few other insects can withstand the large hydrodynamic forces, thereby decreasing competition and predation [6–8]. Some blepharicerid larvae attach so strongly that forces greater than 600 times their body weight are needed to detach them perpendicularly from the rocks^a^, surpassing in attachment strength all other insects known to date [9, 10].

**Figure 1.**
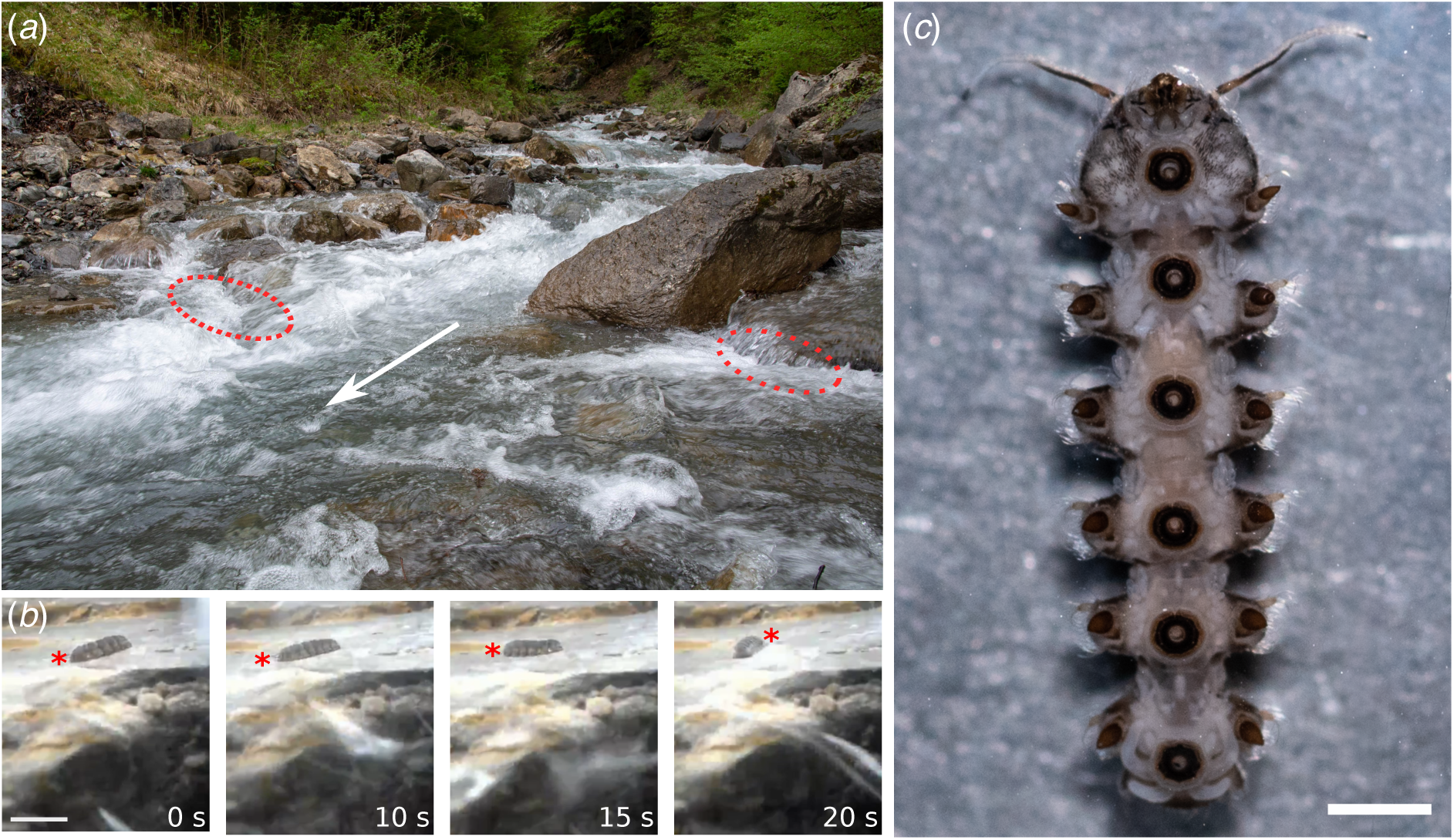
Habitat and crawling locomotion of *Liponeura cinerascens* larvae. (*a*) Fast-flowing alpine river (Oltschibach, Switzerland), one of the collection sites for *L. cinerascens*. Encircled areas indicate sampling locations. (*b*) Selected frames from *in situ* underwater video recording of *L. cinerascens*, showing the larva (head marked with an asterisk) performing a clockwise turn to crawl against the flow (arrowhead points downstream). See supplementary materials SV1 for full video. Scale bar 10 mm. (*c*) Ventral view of 4^th^ instar *L. cinerascens* larva collected from the study site shown in (*a*). Scale bar 1 mm.

Previous studies have investigated how net-winged midge larvae manage to attach, forage, and grow in their natural habitats [1, 3, 5, 7, 11–19]. The functional morphology and development of blepharicerid larvae was reviewed and studied in detail by Rietschel over 50 years ago [11]. The larvae have streamlined flat bodies to minimise drag and, unlike most other rheophilic invertebrates that use hooks and claws to attach to biofilm-covered rocks [20, 21], these larvae rely primarily on suction: each larva has six complex ventromedian suckers with central pistons that are controlled by strong muscles, similar to stalked suckers in decapods [22]. The cavity that houses the piston (called “Manschette” (cuff) by Rietschel and “Chitinsäckchen” (cuticle sac) by other authors [3, 23], translated from German^b^) has a thick sclerotised wall, presumably to withstand the low pressures generated when the piston is raised for suction attachment [11]. The suction disc, which is the ventralmost component of the suction organ, comes into contact with the surface and serves as the main point of attachment. A seal (closure of the suction chamber to minimise leakage of water into the contact zone) is required for suction-based attachments, biological or synthetic. Therefore, it has been suggested that in net-winged midge larvae, the seal is provided by the fringe layer, a thin flexible membrane with hairy projections around the perimeter of the disc, which moulds to the surface contours upon contact [11, 24]. Rietschel also hypothesised that in order to increase shear resistance against the strong hydrodynamic drag forces in fast-flowing waters, the larva uses small cuticular hair-like projections (called microtrichia^c^) as “anchors” to penetrate into the biofilm layer [11]. This idea, however, has yet to be tested experimentally.

A conspicuous V-shaped opening at the anterior side of the suction disc, called a V-notch, has led to a number of different interpretations in previous studies. Rietschel proposed that water within the suction organ is expelled through this V-notch when the disc forms a new attachment to the surface [11], while Frutiger believed that the structure serves both as a water outlet and the initiation point for detachment [24]. While Hora also thought that the V-notch served as an outlet, he went further to propose that the membranes around the V-notch could function like a backwater valve or a heart valve where fluid is pushed out but prevented from flowing back into the contact zone [8]. There was, however, no detailed characterisation of the V-notch structure or the surrounding membranes to support this claim; hence, it is still unclear how the larva controls the V-notch to open the seal.

Despite the wealth of knowledge from these earlier light microscopy studies on the blepharicerid suction organs, many questions remain unexplored until now. The advancement of imaging technologies and biomechanical methods has enabled us to probe deeper into how these organs generate effective attachments. Furthermore, there is a growing interest from the engineering and applied sciences to adopt principles from biological suction organs to create synthetic suction devices that can out-perform existing technologies [25–30]. Until now, most of the engineering inspiration has been drawn from two well-studied animals which use suction – the octopus and the remora suckerfish. In this study, we present an updated morphology of the suction organs in blepharicerid larvae (using two species – *Liponeura cinerascens* and *Liponeura cordata*) based on evidence from laser confocal scanning microscopy (LCSM), X-ray computed microtomography (micro-CT), and scanning electron microscopy (SEM). Furthermore, we present for the first time *in vivo* observations of the contact zone of suction organs using interference reflection microscopy (IRM).

## 5. Results

The attachment organs of Blepharicid larvae resemble in shape the suction organs from octopus and squid (Fig. 2*a*). The blepharicerid suction organ consists of the suction disc (Fig. 2*b, c*), which contacts the surface and has a central opening that connects to the suction chamber. The suction chamber is stabilised by a ring-shaped cuticular cuff and can be evacuated by a central piston, which has a cone-shaped cap. The cuff is surrounded by a circular cuticular fold (Fig. 2*a*), which has muscles inserting into it and is likely to help move or position the suction organ (Fig. 2*b, c*).

**Figure 2.**
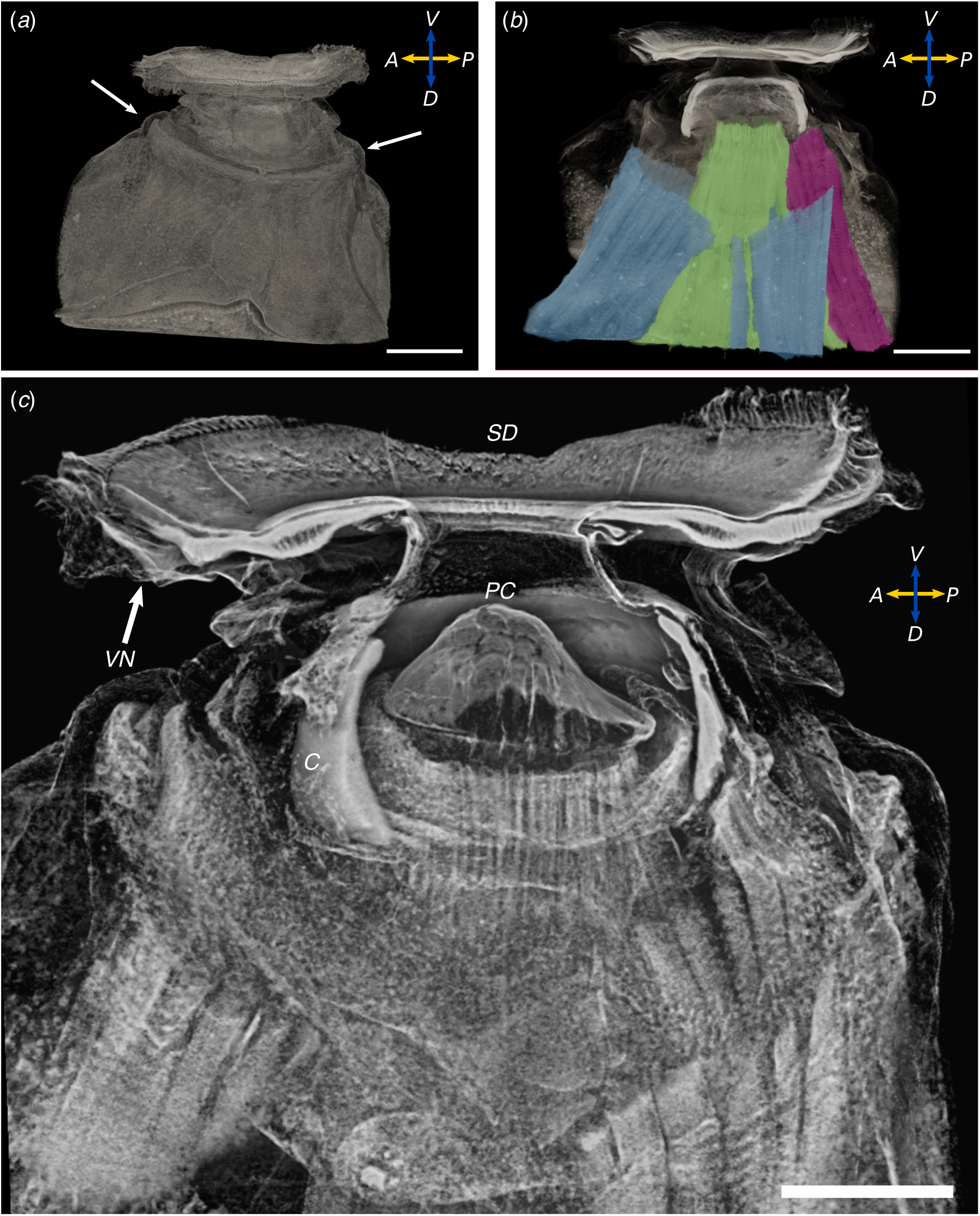
Suction attachment organ from *Liponeura cordata*; 3D reconstruction from micro-CT data. (*a*) External morphology of the organ (arrows point to the cuticular fold). (*b*) Sagittal plane showing three muscle groups inserting into the suction organ (green: piston muscles, magenta: cuff muscles, blue: cuticular fold muscles). (*c*) Sagittal plane highlighting the suction disc (SD), V-notch (VN), piston cone (PC), and the cuff (C). Major muscle groups are also visible (for labels refer to *b*). A: anterior, *P*: posterior, *V*: ventral, *D*: dorsal. Scale bars 100 *µ*m.

### 5.1. Suction disc

The suction disc is the ventralmost part of the suction organ and is therefore the structure that contacts the surface (Fig.2*c*). Suction disc diameters for fourth instar larvae of *L. cinerascens* and *L. cordata* were 426 ± 11 *µ*m (n = 3; average ± 1 standard deviation hereafter) and 377 ± 9 *µ*m (n = 2), respectively. Morphometrics of suction disc diameters and larval body lengths of several species of blepharicerids have been reported elsewhere [24] and are in good agreement with our values. The disc appears darkly pigmented and is stabilised by sclerotised internal radial rays [11]. Various functionally-important structural adaptations of the disc have been revealed through interference reflection microscopy (IRM), SEM, confocal microscopy, and micro-CT techniques, and are described in detail in the following sections.

#### 5.1.1. Microstructures on the suction disc

*In vivo* observations of larvae attaching to a glass coverslip using IRM, and SEM images of the surface of suction organs showed various microstructures present on the disc (Fig. 3). Beginning with the most peripheral structures in *L. cordata*, two rings of hairs called the “fringe” by previous authors form the outer margin of the suction disc [11, 24]. Some fringe hairs came into close contact with the surface, but these did not form a continuous sealing rim around the circumference. Instead, we observed a region just proximal to the fringe forming an almost continuous ring of close contact with the surface (as evident from the dark zero-order interference; Fig. 3*a*). It is likely that this zone is important for sealing the suction chamber to allow the suction organ to generate and maintain sub-ambient pressures. Based on the *in vivo* IRM images, the width of the zone in close contact was 1.9 ± 0.01 *µ*m (n = 2 larvae; see Table 1 and Fig. 3*a*). SEM images of this region showed a pronounced cuticular rim of the same width (1.9 ± 0.05 *µ*m, n = 2 larvae) at this location, formed by short flat cuticular projections joined side-by-side in an uninterrupted regular series (Fig. 3*b, c*). Each structure, referred to as a rim microtrichium, was 0.64 ± 0.13 *µ*m wide (n = 4 larvae; see Table 1) and oriented approximately parallel to the surface (Fig. 3*c*). These rim microtrichia repeated all the way around the perimeter of the disc, forming a continuous region of close contact around the attachment organ. In contrast, we did not observe such rim microtrichia around the disc rim perimeter in *L. cinerascens* (Fig. 3*d*). Instead, SEM images of the suction discs showed a dense array of short and upright rim microtrichia at the same location as the rim microtrichia in *L. cordata* (Fig. 3*e, f*).

**Table 1.**
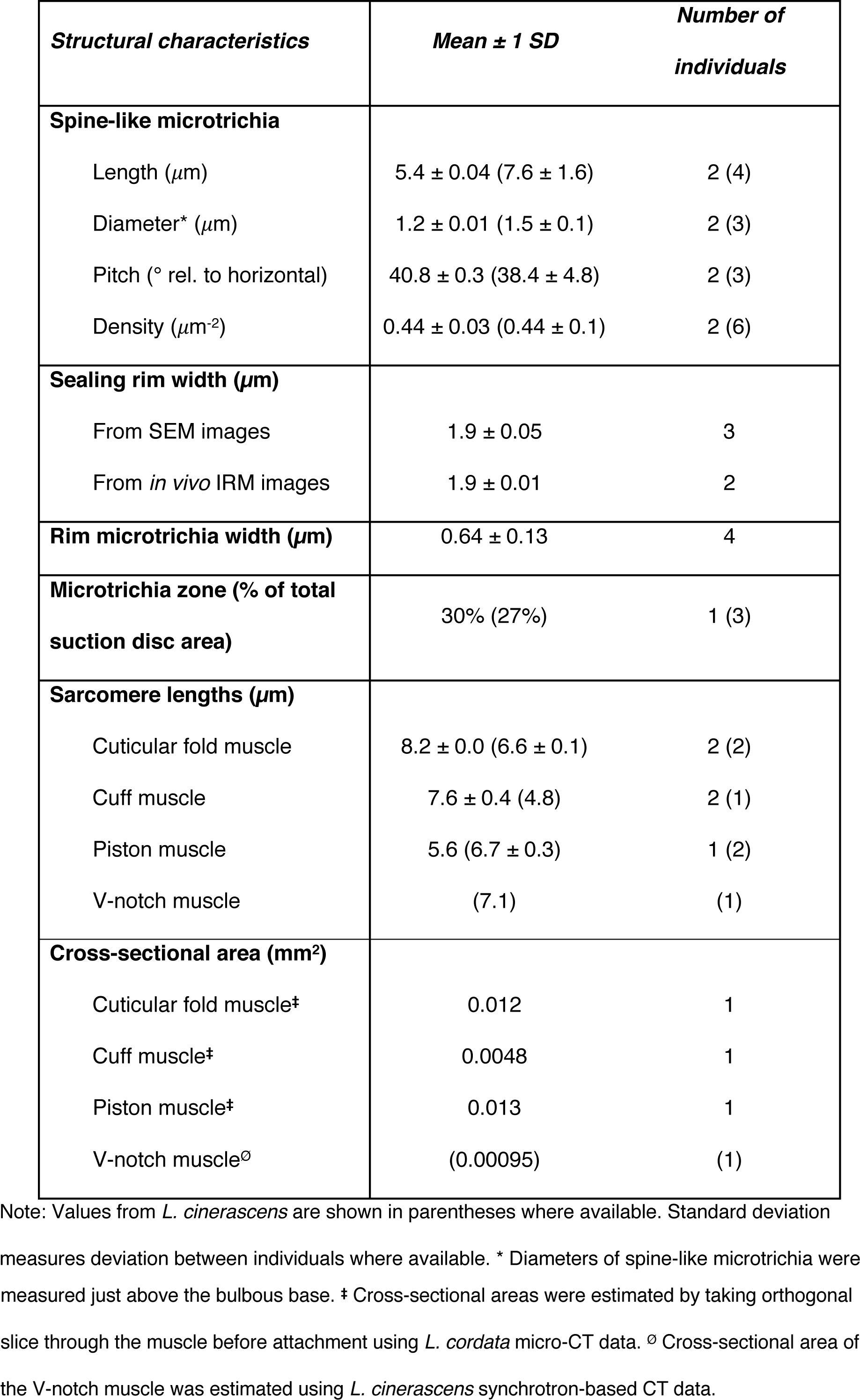
Summary of measurements of relevant structures from *Liponeura cordata* suction organs.

| **Structural characteristics** | **Mean ± 1 SD** | **Number of individuals** |
| --- | --- | --- |
| **Spine-like microtrichia** |  |  |
| Length ($\mu\text{m}$) | 5.4 ± 0.04 (7.6 ± 1.6) | 2 (4) |
| Diameter* ($\mu\text{m}$) | 1.2 ± 0.01 (1.5 ± 0.1) | 2 (3) |
| Pitch ($^{\circ}$ rel. to horizontal) | 40.8 ± 0.3 (38.4 ± 4.8) | 2 (3) |
| Density ($\mu\text{m}^{-2}$) | 0.44 ± 0.03 (0.44 ± 0.1) | 2 (6) |
| **Sealing rim width ($\mu\text{m}$)** |  |  |
| From SEM images | 1.9 ± 0.05 | 3 |
| From *in vivo* IRM images | 1.9 ± 0.01 | 2 |
| **Rim microtrichia width ($\mu\text{m}$)** | 0.64 ± 0.13 | 4 |
| **Microtrichia zone (% of total suction disc area)** | 30% (27%) | 1 (3) |
| **Sarcomere lengths ($\mu\text{m}$)** |  |  |
| Cuticular fold muscle | 8.2 ± 0.0 (6.6 ± 0.1) | 2 (2) |
| Cuff muscle | 7.6 ± 0.4 (4.8) | 2 (1) |
| Piston muscle | 5.6 (6.7 ± 0.3) | 1 (2) |
| V-notch muscle | (7.1) | (1) |
| **Cross-sectional area ($\text{mm}^2$)** |  |  |
| Cuticular fold muscle<sup>‡</sup> | 0.012 | 1 |
| Cuff muscle<sup>‡</sup> | 0.0048 | 1 |
| Piston muscle<sup>‡</sup> | 0.013 | 1 |
| V-notch muscle<sup>⊙</sup> | (0.00095) | (1) |

Note: Values from *L. cinerascens* are shown in parentheses where available. Standard deviation measures deviation between individuals where available. \* Diameters of spine-like microtrichia were
measured just above the bulbous base. ‡ Cross-sectional areas were estimated by taking orthogonal slice through the muscle before attachment using *L. cordata* micro-CT data. ∅ Cross-sectional area of the V-notch muscle was estimated using *L. cinerascens* synchrotron-based CT data.

**Figure 3.**
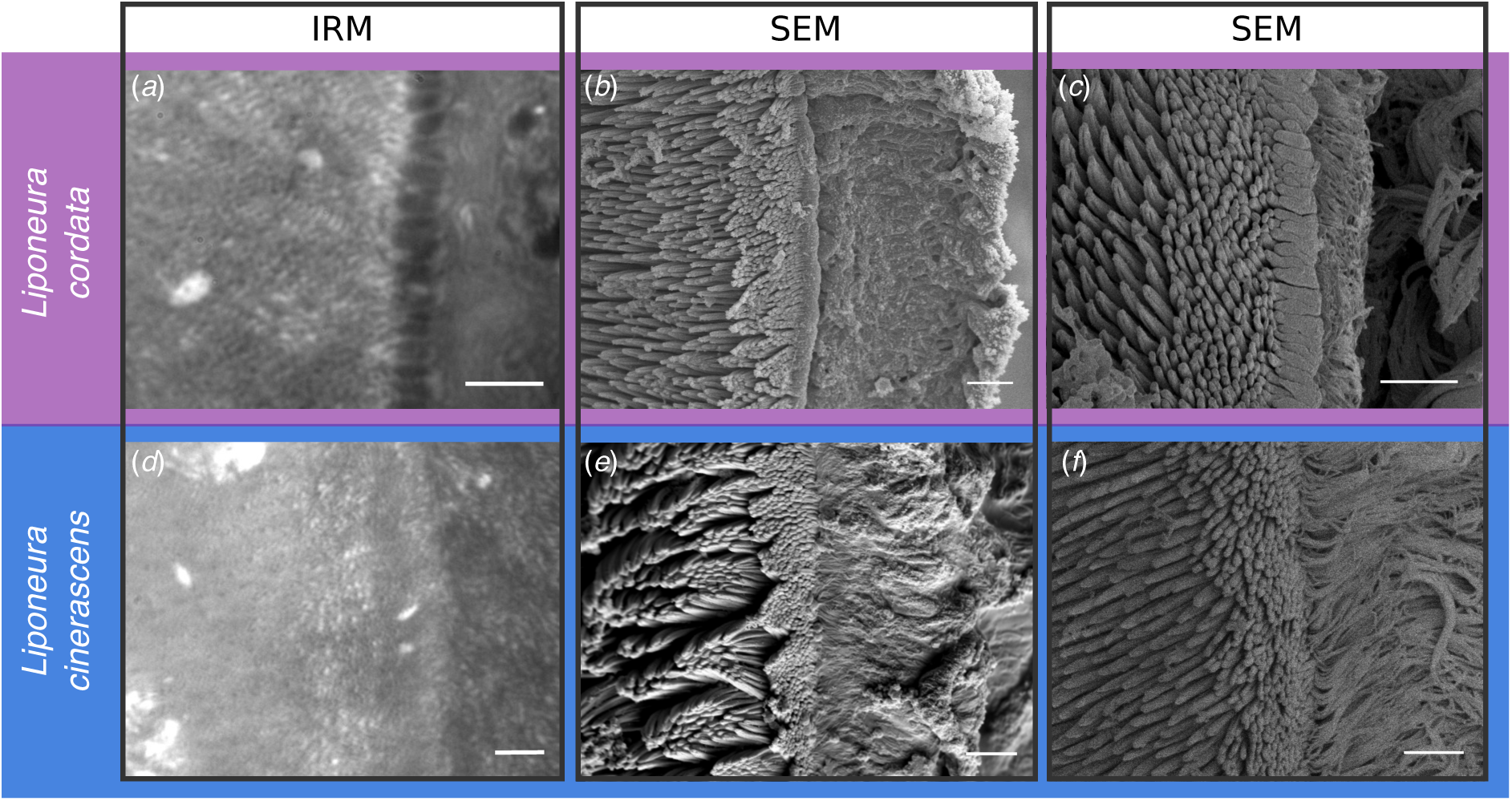
Microstructures on the suction discs of two *Liponeura* species. In *L. cordata.* (*a* – *c*), there is a well-defined contiguous ring of rim microtrichia. (*a*) *in vivo* image of a suction disc in contact with glass using IRM shows a row of rim microtrichia in close contact with the surface (black). (*b, c*) SEM images of the rim highlight the distinct flattened large microtrichia present in *L. cordata*. in (*c*). *(d*) In comparison, the rim zone is less distinct in *L. cinerascens*. (*e, f*) SEM images highlight that in *L. cinerascens*, a rim is formed by a region of short, upright microtrichia. Note the fringe layer to the right of the rim. Scale bars (*a*) & (*d*): 3 *µ*m, (*b*) & (*e*): 5 *µ*m, (*c*) & (*f*): 3 *µ*m.

The suction disc rim was not the only region of the suction disc that came into close contact with the substrate during attachment. *In vivo* IRM recordings showed a dense array of microstructures immediately proximal to the rim (towards the central opening) also coming into contact (Fig. 4; supplementary video SV2). Individual microstructures formed small, roughly circular contact areas on glass coverslips (0.07 ± 0.2 *µ*m^2^, n = 2 *L. cinerascens* larvae), indicating that the tips made close contact. Video recordings showed a pulsing movement of the microstructures, which may be caused by the flow of water into and out of the contact zone (supplementary video SV3). SEM images of the suction disc show that these microstructures are spine-like microtrichia (Fig 3*b-c, e-f*). These microtrichia were present in both *Liponeura* species and were limited to an outer zone that covers approximately 30% of the total suction disc area (Fig. 5*a*). They are longer than the rim microtrichia and are curved, tapered in shape, and have distinct bulbous bases (Fig. 5*b*). There is a gradual transition from the flat rim microtrichia to the longer spine-like microtrichia. While the morphology of the short rim microtrichia differs between the two *Liponeura* species, the spine-like microtrichia structure is highly conserved (supplementary figure 1). Additional characteristics of the spine-like microtrichia, including their dimensions, density, and pitch (relative to the suction disc plane) are summarised in Table 1.

**Figure 4.**
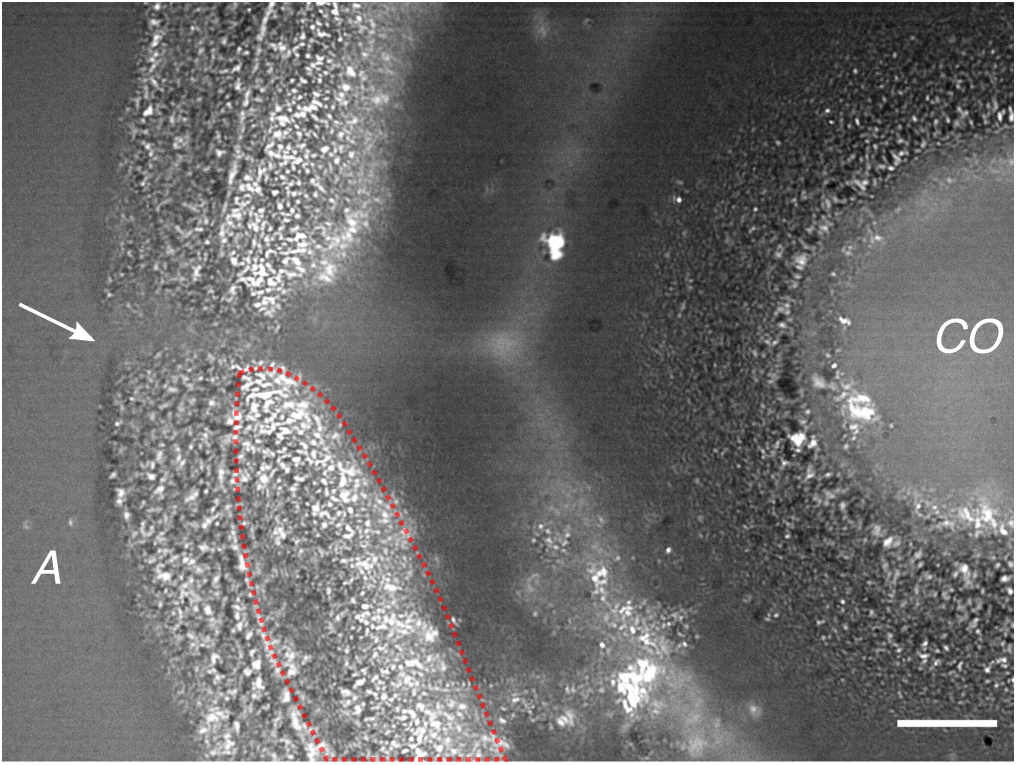
Suction disc of *L. cinerascens* larva in contact with a glass surface, visualised using *in vivo* interference reflection microscopy (IRM). Small, roughly circular black dots represent microtrichia in close contact to the surface (see region encircled in red). White regions are slightly further away, and grey regions even further from the surface. Arrow points to the V-notch (see text for details). *A*: anterior; *CO*: central opening. Scale bar: 15 *µ*m.

**Figure 5.**
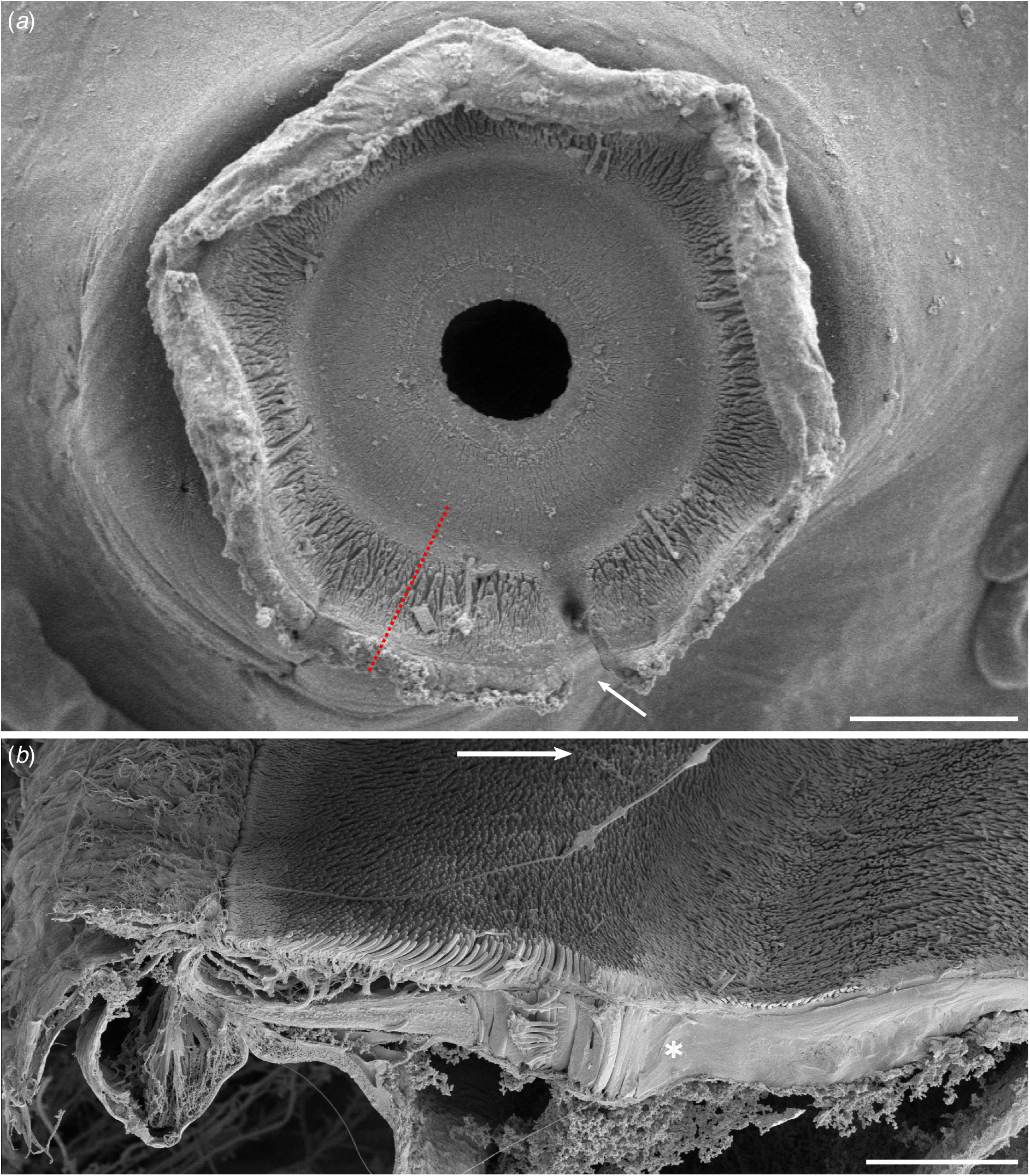
Microtrichia on the ventral surface of a 4^th^ instar *Liponeura cordata* suction disc. (*a*) Cryo-SEM showing the location of the microtrichia region. Note the V-notch is in the open state (see arrow). The red dotted line indicates the approximate length and location of the freeze-fracture represented in *b.* Scale bar 100 *µ*m. (*b*) Freeze-fracture SEM highlighting the different types of microtrichia found on the suction disc. The rim microtrichia are flat and short, while the spine-like microtrichia immediately proximal are long, curved, and oriented towards the centre. The central zone with short and approximately perpendicular microtrichia begins inward from the palisade layer (marked with an asterisk). Arrow points towards the centre of the suction disc. Scale bar 20 *µ*m.

In addition to the rim and spine-like microtrichia, we observed fine structures near the central opening also making contact (Fig. 4). These fine central microtrichia are much shorter than the spine-like microtrichia, and lack the pitch and curvature (Fig. 5*b*). We observed that when the piston was retracted (lowering the internal pressure), the central area was brought closer to the surface and a larger area of the central microtrichia zone came into contact (supplementary video SV4).

#### 5.1.2. V-notch

At the anterior side of each suction disc, the V-notch sharply disrupts the circular outline of the disc in both *Liponeura* species (Fig. 5*a* and Fig. 6). When the V-notch is shut, the sealing rim is almost continuous (Fig. 4), but when open at its widest aperture, the V-notch gap is 2.6 ± 0.3% (n = 3, *L. cinerascens*) of the perimeter of the disc (i.e., for a fourth instar larva with 440 *µ*m disc diameter, the V-notch is around 36 *µ*m wide). The V-notch extended from the edge until the end of the spine-like microtrichia zone (33.3 ± 2.7% of the suction disc radius, from n = 3, *L. cinerascens*) and the area surrounding the base of the V-notch lacked spine-like microtrichia. Through *in vivo* IRM recordings, we clearly observed local, active movements of the V-notch: when the suction disc was firmly attached to a surface, the V-notch was closed, but prior to detachment during locomotion, the two lateral flanks of the V-notch were peeled open starting from the outer end, thereby breaking the seal and rapidly equalising the pressure (Fig. 6*a*, and supplementary materials SV5). We also observed frequent rapid twitching of the flanks of the V-notch, which resulted in quick opening and closing without full detachment. From LCSM images of *L. cinerascens*, we found two long tendon-like cuticular apodemes that insert distally into the flanks of the V-notch and then wrap around the cuff (but within the outer wall of the stalk of the suction disc) to project into the lateral-posterior direction (Fig. 6*b*). Although the ends of the apodemes were beyond the scanning depth of LCSM, we successfully traced the entire length of the structures using 3D renderings from X-ray synchrotron scan data of *L. cinerascens* (Fig. 7). We discovered that each apodeme extends deep into the body, beyond the base of the cuticular fold, where it attaches to a dedicated muscle (Fig. 7*b*). The apodeme was estimated to be 500 - 600 *µ*m long (n=1 larva based on the synchrotron-based data) and 3.1 ± 0.8 *µ*m in diameter (n = 2 *L. cinerascens* larva based on LCSM). The sarcomere length of a V-notch muscle was 7.1 *µ*m (n = 1 larva). The V-notch apodemes did not appear to be attached to the cuff wall or to any surface as they circumvented around the cuff and into the body. The cross-sectional area of the V-notch muscles were smaller than other muscle groups associated with the suction organ (piston, cuff, cuticle fold muscles; see Table 1).

**Figure 6.**
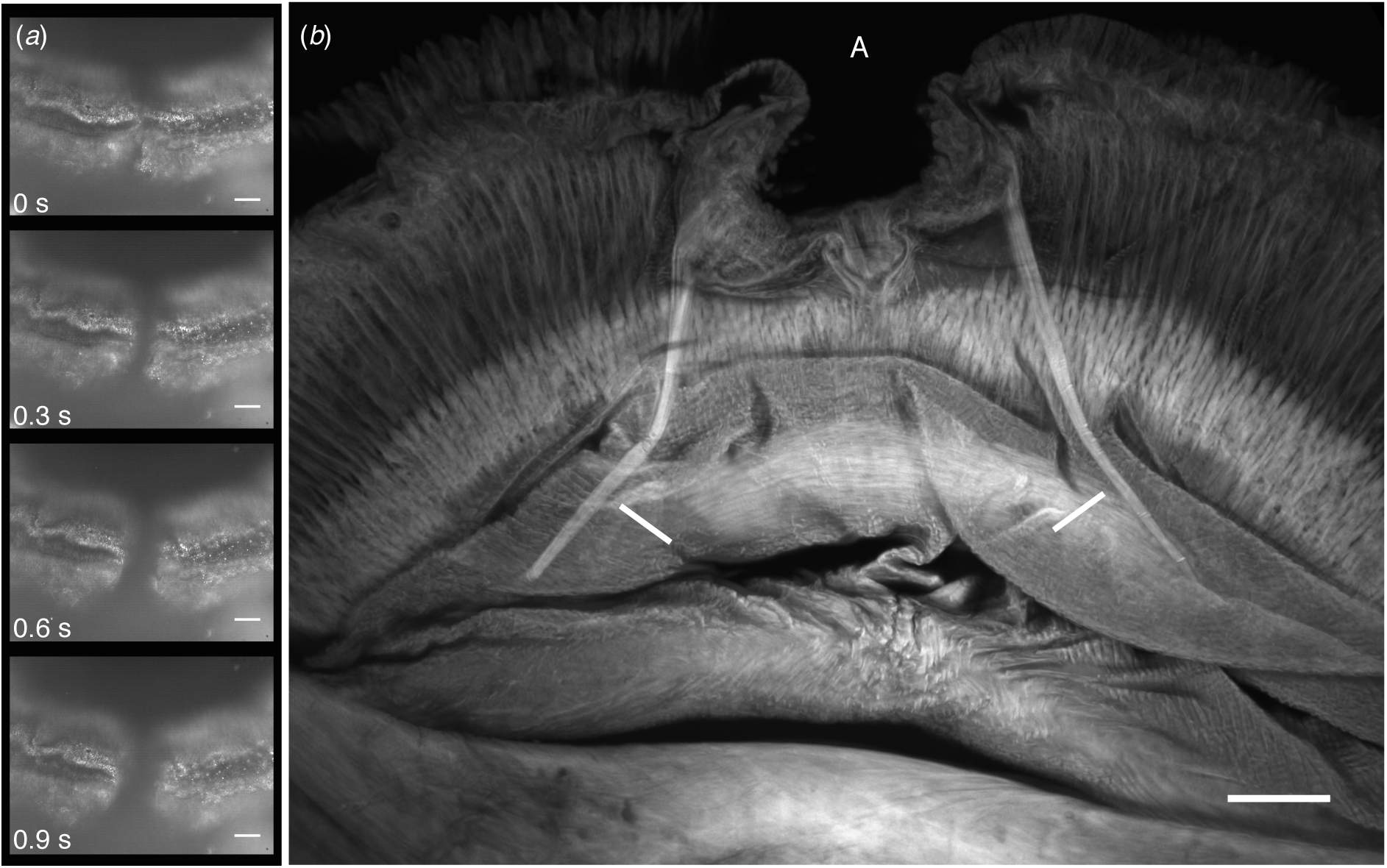
The V-notch system and its movement in a suction organ from *Liponeura cinerascens*. (*a*) Selected frames from *in vivo* IRM recording showing the V-notch being pulled open from the flanks (see supplementary video SV5 for the full video). (*b*) Confocal microscopy image of a Congo Red-stained suction organ reveals the two apodemes attaching to the V-notch flanks (see arrows) that mediate active muscular control of the V-notch. Scale bars 30 *µ*m. *A*: anterior.

**Figure 7.**
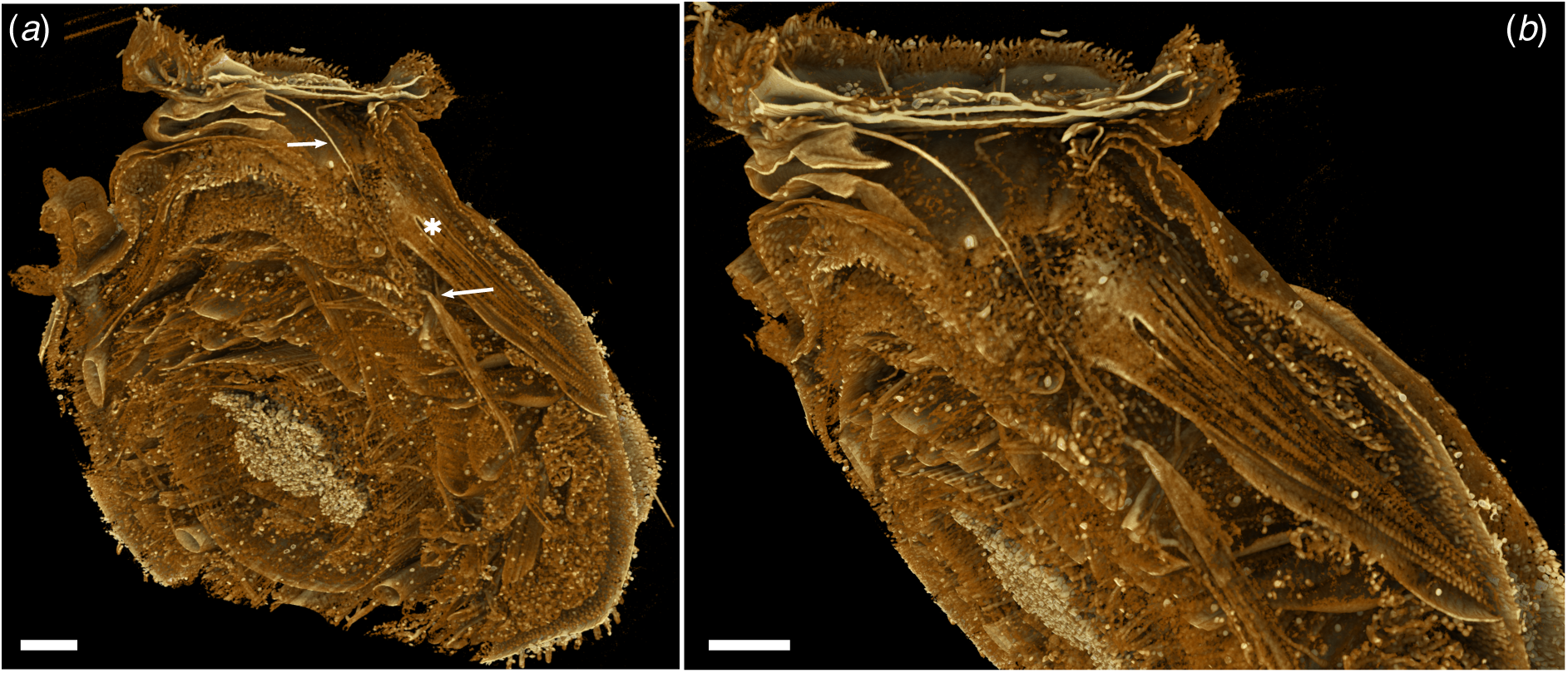
The V-notch apodeme extends posteriorly into the body to attach to a dedicated muscle. 3D reconstruction using X-ray synchrotron data of *Liponeura cinerascens*. (*a*) Overview of the suction organ digitally dissected to reveal the apodeme and the V-notch muscle (arrows). The muscle attaches to the inner dorsal wall. Note the cuff muscles next to the V-notch muscles (marked by an asterisk). (*b*) Lateral view of the apodeme and V-notch muscle. The curvature of the apodeme is clearly visible. Scale bars 100 *µ*m.

In addition to the apodemes and muscles of the V-notch, a second novel feature of the V-notch was identified: in a sagittal view rendering from micro-CT data, it can be seen that the V-notch is shaped like a cupped hand and distinctly juts out further dorsally than the posterior side of the suction disc that has no V-notch (Fig. 8). This membranous structure may have a valve-like function, as it could be seen widening and fluttering as the piston was lowered and water expelled through the valve, but when the piston was raised and the internal pressure lowered, the valve stopped moving (supplementary video SV6). It is likely that this structure and the associated movements serve to seal the V-notch, as proposed previously without direct evidence [8].

**Figure 8.**
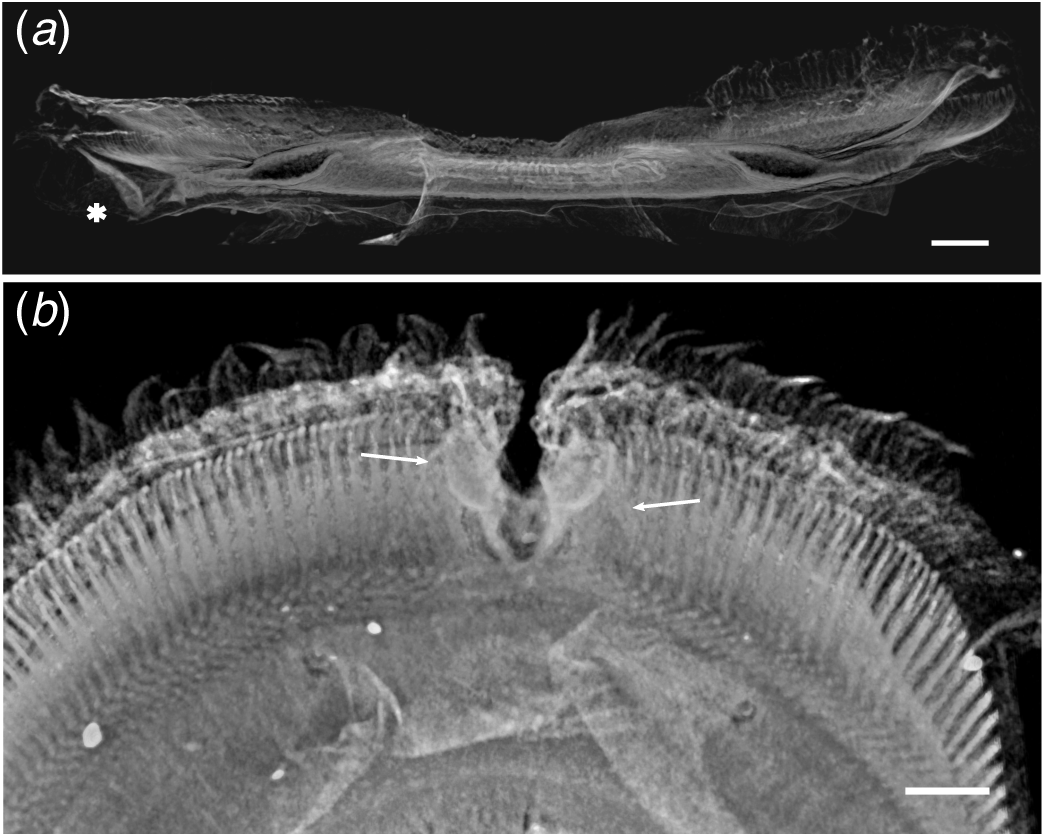
Detailed views of the V-notch valve-like structures from *Liponeura cordata*, reconstructed from micro-CT data. (*a*) Sagittal view of a suction disc rendered to reveal internal structures. The V-notch flap juts out dorsally from the suction disc (marked with an asterisk) compared to the opposite (posterior) side of the disc and could function as a backwater valve to prevent influx of water during piston contraction (see supplementary materials video SV6). (*b*) Dorsal view of the V-notch valves showing their flap-like morphology. Note that the V-notch apodemes attach to the flanks of the flaps (arrows). Scale bars 30 *µ*m.

### 5.2. Piston and cuff

The blepharicerid larva uses a central piston to lower the pressure within the contact zone, similar to the stalked suckers of decapods [22]. The conical cap of the piston is comprised of dark sclerotised cuticle. This cone is surrounded by the cuff, which has similar characteristics and is connected to the cone via a thin flexible membrane. A reservoir of water is always kept within the volume enclosed by the cuff, which is important when generating pressure differentials as water is an incompressible fluid, so that small volume changes from piston movements can lead to large reductions in internal pressure. A pair of large muscles called the piston muscles extend from the left and right sides of the dorsal cuticle and attach to the piston cone via multiple thread-like intracellular attachments, referred to as tonofibrils by Rietschel [11] (Fig. 2*c* and supplementary figure 2). The muscles are positioned such that contraction leads to a movement perpendicular to and away from the surface. Sarcomere lengths from the piston muscles of fourth instar *L. cinerascens* were on average 6.7 ± 0.3 *µ*m (n = 2 larvae) and 5.6 *µ*m for fourth instar *L. cordata* (n = 1 larva).

Besides piston muscles, there is a pair of large muscles that extend from the inner dorsal wall and attach to the base of the membrane surrounding the cuff wall (Fig. 2*b*, magenta). These cuff muscles are found only at the posterior end of the cuff, hence unlike the piston muscles, contraction of cuff muscles leads to an asymmetric directional pull on the organ. Cuff muscle sarcomere lengths from *L. cinerascens* were 4.8 *µ*m (n = 1 larva) and 7.6 ± 0.4 *µ*m *L. cordata* (n = 2).

### 5.3. Cuticular fold

The cuticular fold is formed by generous folding of the outer cuticle (Fig. 2*a*) and partly encapsulates the piston apparatus (Fig. 2*b*, blue). Near the cuticular fold, to the left and right sides of the suction organ, we observed two pairs of large muscles called cuticular fold muscles that could help to manoeuvre the suction organ. This is in agreement with previous studies that illustrated the muscles attaching to the organ such that contraction would lead to a force acting on the entire organ [3, 11]. Note that the cuticular fold muscles attach to the suction organ and not the fold itself so that a contraction would retract the organ into the body of the larva. In *L. cinerascens*, the sarcomeres from the cuticular fold muscles were on average 6.6 ± 0.1 *µ*m in length, whereas in *L. cordata* they were 8.2 ± 0.0 *µ*m (n = 2 for each species). As both muscle pairs are located on the lateral sides of the organ, contraction is likely to produce movements in either vertical or lateral directions and limited movements in forward or reverse.

## 6. Discussion

### 6.1. Increasing shear resistance in suction attachments

The blepharicid larvae have evolved highly complex suction organs with unique structural adaptations for powerful and effective attachment on various surfaces. One such feature is the dense array of microtrichia within the suction disc. Rietschel proposed that these cuticular projections, which he compared to the gecko’s adhesive setae, help the larva resist lateral forces by penetrating into biofilm layers and anchoring the animal [11]. While the suction organs of male diving beetles [31, 32] and medicinal leeches [33] have smooth surfaces, the surfaces of suckers from octopus, remora fish, and clingfish have similar small projections or spine-like structures [30, 34–37]. The proposed primary function of these structures is to come into contact with the surface and increase resistance to shear forces. In the remora fish, stiff mineralised conical structures called spinules are found at the tip of lamellae and can be actively rotated erect or flat using muscles attached to the lamellae [28, 35, 38]. The remora spinules have an aspect ratio of approximately 2:1 (average length and width of roughly 500 *µ*m and 270 *µ*m, respectively) and can interlock with surface asperities using pointed tips angled at 34° relative to the horizon [28]. They are arranged in consecutive rows and their tips face the posterior end of the fish. While the blepharicerid microtrichia are different to the remora spinules in that they are thinner and longer (aspect ratio of around 5:1), more numerous (tens of thousands per individual compared to around a thousand in remora) and arranged as arrays covering the suction disc (compared to row arrangement in remora), they have similar shapes, tips, and pitch (Fig. 9). Since remora spinules and blepharicerid microtrichia are both found on suction attachment organs that need to withstand strong drag forces, the main function of these structures could be similar - to increase friction force on rough substrates and thereby improving attachment performance. Peak attachment force measurements on live remora fish on smooth versus rough (shark-skin) surfaces showed that higher shear forces were needed to detach them from rough surfaces [35], suggesting that the spinules are important in increasing the friction force between the suction organ and the rough surface; subsequent quantitative studies confirmed that remora spinule tip contact significantly enhances friction by interlocking with asperities of rough surfaces [28, 38]. Even on smooth surfaces, small fibres could increase friction by making tip or side contact [39–41], but typically, adhesion-based shear forces are strongly reduced underwater. Overall, these findings support the idea that microtrichia from blepharicerid suction discs increase the resistance to shear forces, which may be critical for animals living in high-drag aquatic environments. Further studies are needed to quantify the friction forces from the microtrichia zone to better understand the function of this characteristic morphology.

**Figure 9.**
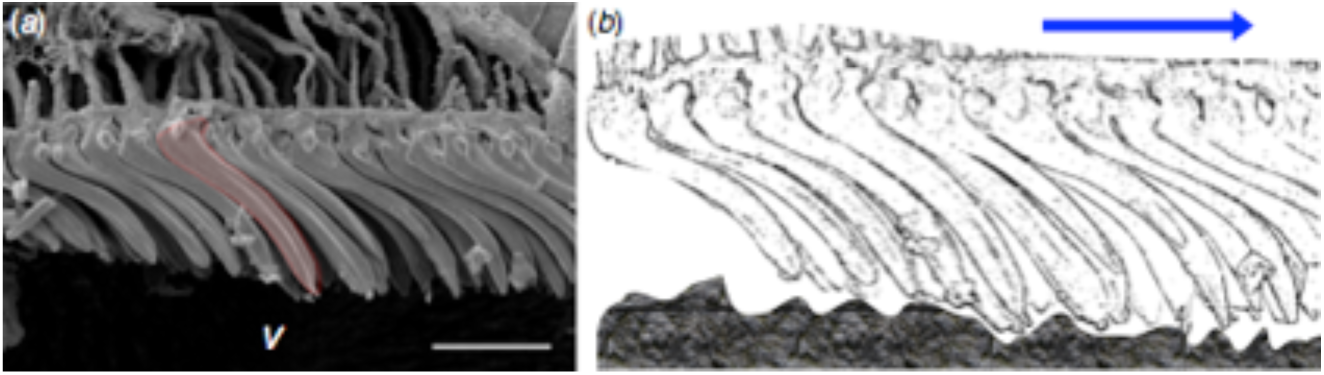
Microtrichia from *Liponeura cinerascens*. (*a*) Freeze-fracture SEM of a suction disc showing a single microtrichium (shaded in red) to highlight the curvature and pitch angle (see Table 1 for measurements). *V*: ventral. Scale bar 5 *µ*m. (*b*) Schematic depiction of microtrichia making tip contact with a hypothetical rough rock surface. Arrow points towards the centre of the suction disc.

### 6.2. Sealing in suction organs

A seal is required for suction-based attachment, where a sustained pressure gradient is critical for maintaining adhesive strength. In the case of the octopus and remora fish, previous authors assumed that a soft fleshy rim moulds to the surface roughness to provide an efficient seal [34, 35]. With abalones and clingfish, it has been proposed that a dense network of micrometre-sized fibrous structures - cilia (incorrectly labelled as “nanofibrils” in [42]) and “microvilli” in abalones and clingfish, respectively - helps the adhesive organ mould extremely well to any surface roughness and thereby generate an effective seal [36, 42]. If such fibres are able to displace the water film and form dry adhesive contacts with the surface, they could contribute significantly to both normal and shear detachment forces [36, 42]. The dense array of microtrichia on the surface of blepharicerid suction discs could serve similar functions as the fibrils in clingfish and abalones. It is noteworthy that we found two different rim morphologies: flat rim microtrichia in *L. cordata* (Fig. 3*c*), as opposed to a densely packed array of short rim microtrichia in *L. cinerascens* (Fig. 3*f*). Even without a continuous rim contact as seen in *L. cordata*, it is possible that the dense array of rim microtrichia in *L. cinerascens* can reduce the flow of water into the suction chamber by decreasing the gaps between individual elements to extremely small sizes. As the flow-rate through a narrow slit is proportional to the cube of its width, gap size has a dramatic effect on flow rate. Reduction in flow rate may also apply to the spine-like microtrichia zone, albeit to a lesser degree due to the wider gaps between each fibre. The hierarchy of extremely dense rim microtrichia to dense spine-like microtrichia, however, may help the blepharicerid suction organ generate a good seal even on natural substrates with a wide range of surface roughness length scales.

### 6.3. Generating large pressure gradients using specialised piston and musculature

The attachment strength of the blepharicerid suction organ is directly related to the pressure difference between the organ and the external environment. Upon contraction, the piston muscles raise the piston away from the surface, thereby significantly lowering the internal pressure relative to the surroundings (assuming there is close to zero leakage), and leading to a powerful suction force. As water is nearly incompressible, the pull will increase the internal volume only minimally but may elastically deform the walls of the chamber, and the cuff in particular. The piston muscles must be capable of generating large forces to maximise the potential attachment strengths. Gronenberg *et al.* found that muscle fibres in ant mandibles with long sarcomeres (approximately 5 *µ*m in length) were characteristic of slow and forceful muscles, while short sarcomeres (approximately 2 *µ*m) were typically attributes of fast muscles [43]. Since the average sarcomere lengths of piston muscle fibres were 5.6 and 6.7 *µ*m in *L. cordata* and *L. cinerascens*, respectively, the piston muscles may also be adapted for high force as opposed to speed. Furthermore, we found that the cross-sectional area of the piston muscles was the largest out of the muscle groups within the suction organ. As a stronger piston pull leads to a larger pressure gradient, and as muscle force scales with the cross-sectional area, the piston muscles are designed to generate strong forces for powerful attachments.

### 6.4. Mechanism for rapid detachment of a suction organ

The V-notch appears to be a unique feature of the blepharicerid suction discs as no such structures have been reported from other well-studied suction organ systems. Previous researchers proposed that the V-notch provides a detachment mechanism for the suction organ [3, 8, 11, 24]. Since these larvae are able to locomote using the suckers (up to 0.96 mm s^-1^ in rapid evasive movement, or approximately 0.1 body lengths per second [7]), they need to repeatedly and rapidly release the powerful suction attachment and detach the suction discs. From video footage of moving larvae on a glass aquarium surface, Frutiger suggested that the detachment begins at the V-notch and spreads radially to create an opening for sudden depressurisation of the organ [24], but the exact sequence of this process was still unclear (Fig.5 in [24]). Our recordings of larval locomotion on glass showed that when the piston muscle was contracted, and the suction organ was fully engaged, the V-notch remained closed so that the surface contact of the rim of the suction disc was continuous (supplementary video SV6). As the larva prepared for detachment, the V-notch apodemes were pulled, peeling the V-notch flanks away from the surface. Subsequent influx of water through this opening equalised the pressure within the suction chamber so that the suction disc could be lifted from the substrate, presumably using the strong cuticular fold muscles. We observed fine control of the V-notch, including independent movements of each V-notch flank, and twitching of the flanks; on several occasions, one flank was pulled just a few micrometres before relaxing and closing the gap. Such movements are unlikely to be passive (i.e., caused by external flow, or displacements from larger-scale muscle-driven body movements) given the independent control of each V-notch flank and the speed of the twitching. Indeed, we discovered that the apodemes extend into the body to connect directly to a pair of dedicated V-notch muscles. The larva, therefore, has specific motor control of the V-notch, allowing them to open it independently at a range of speeds. This V-notch system is, to our knowledge, the first description of a mechanism specifically dedicated to detachment of a biological suction organ. It would be fascinating to explore the development of such a long apodeme running extensively through intracuticular space, and how these structures and the entire suction organ are re-built between each moult.

While the V-notch muscles open the V-notch, it is not clear how the V-notch can be closed and sealed. Previous researchers have used the term “valvular gateway” to describe the V-notch structure [8, 16], and Hora further proposed that the membranous flaps of the V-notch are arranged to allow water expulsion but not influx. Our *in vivo* observations provide additional support for this idea, as the membranous flaps could be seen to widen and vibrate from water being expelled as the piston was lowered, while the valves narrowed and ceased to move when the piston was raised and the internal pressure lowered. The valves could be closed passively by the pressure gradient during suction cup attachment. Hence, the V-notch valves could function similar to heart valves, which rely on pressure gradients to open and close during heart contractions to prevent backflow between the atrium and the ventricle [44]. Interestingly, the mitral heart valves also have chord-like attachments called the chordae tendineae that resemble the V-notch and its apodemes [45]. The chordae tendineae primarily act to prevent valvular prolapse by tethering them to the papillary muscles of the inner ventricular wall when the pressure rises from ventricular contraction. The V-notch tendons, in addition to actively peeling open the structure, could also serve as a tether to prevent the V-notch valves from folding into the contact zone when the organ is engaged; we never observed this in our *in vivo* studies.

## 7. Conclusion

The larvae of Blephariceridae need to withstand powerful hydrodynamic forces in their habitats. They rely on six complex suction organs with several unique adaptations to generate strong attachment yet rapid detachment. Firstly, a continuous sealing rim may minimise leakage; it consisted of flat rim elements in *Liponeura cordata*, and a dense array of short microtrichia in *Liponeura cinerascens.* Secondly, a dense region of cuticular projections (microtrichia) probably increases shear resistance on rough surfaces, similar to structures in the suction pad of remora fish helping them to attach onto rough shark skins. Lastly, the V-notch on the anterior side of the suction disc can be actively pulled open via two apodemes attached to its flanks, independently controlled by dedicated V-notch muscles. Despite these new insights into the functional morphology of the blepharicerid suction system, many questions remain open for future investigations. For example, while we have identified four muscle groups, the V-notch, piston, cuff, and cuticular fold muscles, their individual role in manoeuvring the organ and generating the suction pressure still needs to be clarified. Further work is also needed to elucidate the contribution of microtrichia to shear force resistance. A better understanding of these sophisticated natural suction systems could lead to new bio-inspired strategies to attach strongly to smooth and rough surfaces in both wet and dry conditions.

## 8. Methods

### 8.1. Sample collection and maintenance

Specimens were collected from alpine streams in Austria (near Vols, Innsbruck) and Switzerland (near Interlaken). Some of the specimens were immediately fixed in 70% ethanol, while the rest were brought back to the laboratory in Cambridge and maintained in a climate-controlled aquarium tank at 5°C. Rocks collected from the sampling sites provided diatoms and other biofilm as a source of food, and a timer-controlled LED light promoted biofilm growth. Collected larvae were identified to species and developmental stage based on identification keys from Frutiger [12] and Zwick [46]. Due to the difficulty in maintaining the larvae in laboratory conditions, all experiments were conducted within five days of collection.

### 8.2. In vivo imaging using interference reflection microscopy (IRM)

We imaged the surface contact of live blepharicerid larvae with Interference Reflection Microscopy (IRM). The application of IRM to characterize adhesive systems has been described in previous studies on insects and tree frogs [47, 48]. Briefly, an individual larva was placed on a glass coverslip with a small droplet of aquarium water. Excess water was wicked away (a small amount was left to prevent desiccation), and the attached suction organs were observed under green light (546 nm wavelength) using either a 20x or 100x Leica oil immersion objective. Images and videos were recorded using a CMOS USB3 camera (DMK 23UP1300, The Imaging Source Europe GmbH, Bremen, Germany) and their proprietary software (IC Capture v2.4.642.2631). Recordings were subsequently analysed using Fiji [49] (https://imagej.net/Fiji).

### 8.3. Scanning electron microscopy (SEM) of suction organs

Two types of SEM were used to characterise the morphology of the suction organ: a field emission SEM (FEI Verios 460) at the Cambridge Advanced Imaging Centre (CAIC), and a cryo-SEM (Zeiss EVO HD15 with Quorum PP3010T preparation system) at Sainsbury Laboratory Cambridge University. For field emission SEM (FESEM), samples were prepared as follows: Specimens in 70% ethanol were plunge-frozen in liquid ethane then freeze-dried overnight (Quorum Emitech K775X). To visualise internal structures and cross-sectional slices of the suction organs, the samples were plunge-frozen as described, mounted on a custom-designed aluminium block cooled with liquid nitrogen, then fractured using a double-edged razor blade. All samples were mounted on aluminium SEM stubs using double-sided carbon tapes and conductive silver paint to minimise charging. Samples were consequently sputter-coated using 16 to 32 nm of iridium depending on the amount of expected charging. For cryo-SEM, samples were removed from 70% ethanol, mounted on stubs using colloidal carbon glue, then plunged into a pre-cooled Quorum PP3010T preparation system. Frozen fully hydrated samples were coated with 5 nm platinum immediately prior to imaging.

Post-processing of the SEM images, including quantification of the different structures present on the suction disc, was carried out using Fiji [49] (https://imagej.net/Fiji). Figures were produced using GNU Image Manipulation Program (GIMP v2.10.4, https://www.gimp.org/) and Inkscape (Inkscape v0.91, https://inkscape.org/).

### 8.4. X-ray microtomography (micro-CT) of dissected suction organs

A single suction organ from a *Liponeura cordata* fourth instar larva (fixed in 70% ethanol) was dissected and stained in Lugol’s iodine (PRO.LAB Diagnostics, UK) for 4 days. The stained sample was rinsed in 70% ethanol, transferred in ethanol to a specimen holder made from a micropipette tip attached to a needle holder, and sealed with hot-melt adhesive to avoid evaporation. The sample was analysed via X-ray Microscopy (XRM) using a lab-based Zeiss Xradia Versa 520 (Carl Zeiss XRM, Pleasanton, CA, USA) X-ray Microscope, using a CCD detector system with scintillator-coupled visible light optics, and tungsten transmission target. Multiple scans at varying resolution and field-of-view were carried out to give both an overall visualization, and to reveal regions of interest at higher resolution. For the overall scan an X-ray tube voltage of 70 kV, and a tube current of 85 *µ*A were used, with an exposure of between 7000 - 10000 ms, and a total of 3201 projections. An objective lens giving an optical magnification of 20x was selected with binning set to 2, producing an isotropic voxel (3-D pixel) sizes of between 0.5402 – 0.7111 *µ*m. The tomograms were reconstructed from 2-D projections using a Zeiss commercial software package (XMReconstructor, Carl Zeiss), a cone-beam reconstruction algorithm based on filtered back-projection. XMReconstructor was also used to produce 2-D grey scale slices for subsequent analysis. 3D volume rendering and segmentations were carried out using Dragonfly v3.6 (Object Research Systems Inc, Montreal, Canada) and Drishti v2.6.4 [50].

### 8.5. Synchrotron X-ray micro-tomography of dissected suction organs

Microtomographic scans of an ethanol-fixed dissected suction cup was performed at the UFO imaging station of the KIT light source. A parallel polychromatic X-ray beam produced by a 1.5 T bending magnet was spectrally filtered by 0.5 mm Al at a peak at about 15 keV, and a full-width at half maximum bandwidth of about 10 keV. A fast indirect detector system consisting of a 12 *µ*m LSO:Tb scintillator [51], diffraction limited optical microscope (Optique Peter) and 12bit pco.dimax high speed camera with 2016 x 2016 pixels resolution [52] was employed for taking 3,000 projections at 70 fps and an optical magnification of 10X (1.22 *µ*m effective pixel size). The control system concert [53] was used for automated data acquisition. Tomographic reconstruction was performed with a GPU-accelerated filtered back projection algorithm implemented in the UFO software framework40. 3D volume rendering of tomographic data was done using Drishti v2.6.4 [50].

### 8.6. Laser confocal scanning microscopy (LCSM) of dissected suction organs

In order to prepare samples of optimal thickness, each suction organ was dissected from samples fixed in 70% ethanol. For ventral views of the suction disc, excess tissue was further excised to leave only the disc material for imaging. Dissected samples were rehydrated to 1x phosphate buffered solution (PBS) by being placed inside 50% ethanol-50% PBS, 25% ethanol-75% PBS, and 100% 1x PBS for 5 minutes at each gradient. Two different stains were used: Calcofluor white (Fluorescent Brightener 28, Sigma F3543), a non-specific fluorochrome that binds to chitin, cellulose, and other β-1-3 and β-1-4 polysaccharides [54], and Congo Red (Sigma C6277), also a non-specific fluorochrome that has been used to successfully stain small arthropods for LCSM [55].

Calcofluor white (CFW) stock solution (1% w/v) was prepared by dissolving the powder in 1x PBS in a water bath set to 65°C for 15 – 20 minutes. Samples were stained in 0.1% CFW for 4 hours, washed 3 times in 1xPBS, then mounted on glass slides using the method described by Michels [55] in Fluoromount (Sigma F4680).

Congo Red was prepared as described previously [55], with slight modifications: The solution was gently heated in a water bath at 65°C to fully dissolve the powder, and the stock solution was shielded from light using aluminium foil and kept in a dark cupboard. Samples were stained for 24 hours, removed from stain, rinsed 3 times in PBS at 10 minutes per rinse, then mounted as described above.

LCSM images were acquired using Olympus Fluoview FV3000 (Olympus Corp., Tokyo, Japan) using 405 nm excitation wavelength for CFW and 561 nm for Congo Red. Detection wavelengths were 412 – 512 nm and 580 - 680 nm for CFW and Congo Red respectively. Since Congo Red is sensitive to bleaching, image acquisition settings were adjusted accordingly, and stained samples were imaged within 8 hours of mounting. Mounted CFW samples were sufficiently stable and could be imaged over 3-4 days if kept in the fridge. Images were adjusted for contrast, brightness, and false-colouring using Fiji [49] (https://imagej.net/Fiji).

## 9. Declarations

### Acknowledgements

V.K. would like to thank K.H. Muller and J.N. Skepper at the Cambridge Advanced Imaging Centre for their help in preparing and imaging SEM samples. We are grateful to R. Wrightman for his assistance with imaging cryo-SEM samples. We acknowledge the KIT light source for provision of instruments at their beamlines and we would like to thank the Institute for Beam Physics and Technology (IBPT) for the operation of the storage ring, the Karlsruhe Research Accelerator (KARA).

## Ethics approval and consent to participate

Not applicable.

## Consent for publication

Not applicable.

## Availability of data and material

The datasets generated and/or analysed during the current study are not publicly available due to their large file sizes or because of they are used in on-going research but can be made available from the corresponding author on reasonable request.

## Authors’ contributions

V.K. designed, recorded, and analysed data from IRM, SEM, and LCSM, and wrote the paper. R.J. and V.K. collected and analysed the micro-CT data. T.v.d.K., T.F. and V.K. collected and analysed synchrotron microtomography data. W.F. helped design the study, record and analyse IRM data, and provided feedback for the paper. All authors read and approved the final manuscript.

## 10. Supplementary materials

File name: Supplementary figure 1

File format: PDF

Description: Freeze-fracture SEM image of a *Liponeura cinerascens* suction disc showing spine-like microtrichia. The spine-like microtrichia are similar in shape to those found in *L. cordata* (see Fig. 5*b*). Arrow points towards the centre. Scale bar 5 *µ*m.

File name: Supplementary figure 2

File format: PDF

Description: Thread-like attachments between piston muscles and the piston cone. From Liponeura cordata micro-CT data. (*a*) Lateral view of piston cone shows a void volume where the thread-like attachments connect the ends of the muscles to the top of the piston cone. Arrows indicate thread-like attachments. A: anterior, P: posterior, L: left, R: right. Scale bar 50 *µ*m. (*b*) 3D rendering highlighting the thin fibrous nature of the thread-like attachments.

File name: Supplementary video SV1

File format: .mp4

Description: Video of *Liponeura cinerascens* larva crawling on a rock in a torrential alpine river.

File name: Supplementary video SV2

File format: .mp4

Description: Interference reflection microscopy (IRM) video recording of a live *L. cordata* larva attaching to a surface. Scale bar 15 *µ*m.

File name: Supplementary video SV3

File format: .mp4

Description: Interference reflection microscopy (IRM) video recording of a live *L. cinerascens* larva. A suction disc is in contact with the surface and pulsing of the suction disc microstructures are visible. Scale bar 20 *µ*m.

File name: Supplementary video SV4

File format: .mp4

Description: Interference reflection microscopy (IRM) video recording of a live *L. cinerascens* larva attaching to a surface. When the piston is raised, the pressure inside the suction disc is lowered, and a larger portion of the disc is brought into contact. Scale bar 50 *µ*m.

File name: Supplementary video SV5

File format: .mp4

Description: Interference reflection microscopy (IRM) video recording of a live *L. cinerascens* larva. The V-notch is opened from its flanks as a detachment mechanism for the suction organ. Scale bar 50 *µ*m.

File name: Supplementary video SV6

File format: .avi

Description: Interference reflection microscopy (IRM) video recording of a live *L. cinerascens* larva. The V-notch valves contract and get closer to the surface when the piston contracts and lowers the suction organ pressure. When the piston is relaxed, flickering of the valves indicate water flow. Scale bar 30 *µ*m.

## Footnotes

a Calculation based on force and weight measurements from Frutiger [24] on *Hapalothrix lugubris*.

b Numerous German publications on Blephariceridae morphology (e.g. [3],[18], [23]) appear to use the term “chitin” interchangeably with “cuticle” without experimental evidence that the material is specifically chitin. We adopt the term “cuff” to avoid any erroneous characterization of the underlying material.

c These cuticular projections were called microtrichia by Rietschel [11]. Based on the diagrams of developing suction organs provided in his figures, multiple projections appear to form from single epidermal cells, which fits the definition of microtrichia and hence we continue to use this terminology.

