## Supplementary figures for "Sophisticated suction organs from insects living in raging torrents: Morphology and ultrastructure of the attachment devices of net-winged midge larvae (Diptera: Blephariceridae)"

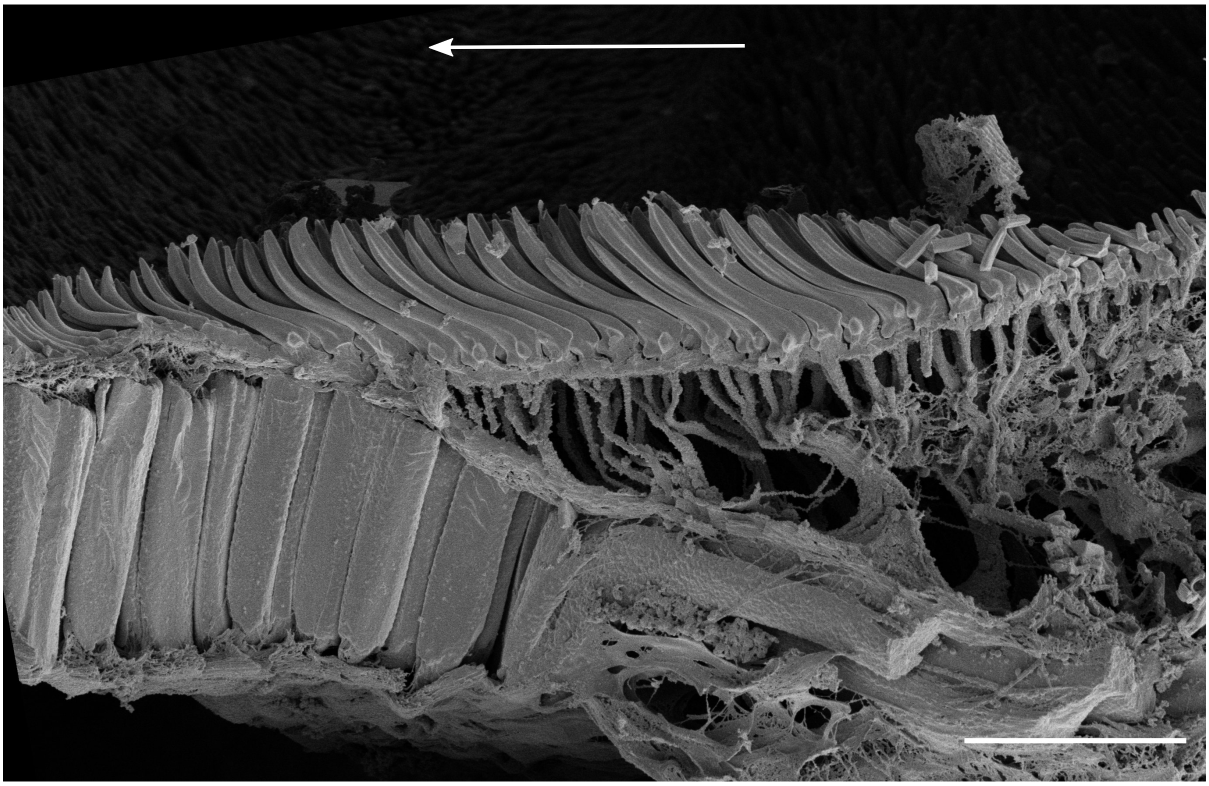

Supplementary Figure 1. Freeze-fracture SEM image of a *Liponeura cinerascens* suction disc showing spine-like microtrichia. The spine-like microtrichia are similar in shape to those found in *L. cordata* (see Fig. 5*b*). Arrow points towards the centre. Scale bar 5 $\mu$m.

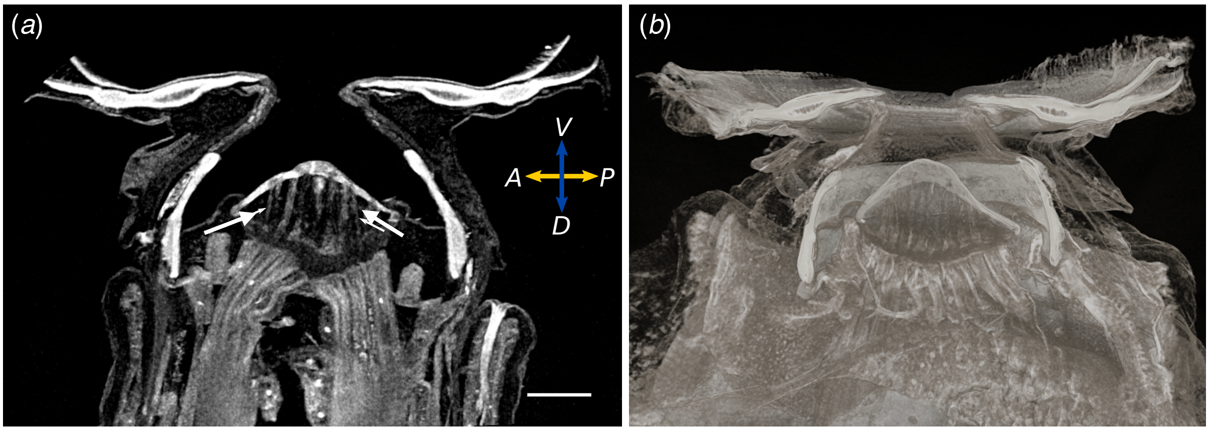


Supplementary Figure 2. Thread-like attachments between piston muscles and the piston cone. From *Liponeura cordata* micro-CT data. (*a*) Lateral view of piston cone shows a void volume where the thread-like attachments connect the ends of the muscles to the top of the piston cone. Arrows indicate thread-like attachments. A: anterior, P: posterior, L: left, R: right. Scale bar 50 $\mu$m. (*b*) 3D rendering highlighting the thin fibrous nature of the thread-like attachments.
